# Genome-scale metabolic reconstruction of the stress-tolerant hybrid yeast *Zygosaccharomyces parabailii*

**DOI:** 10.1101/373621

**Authors:** Marzia Di Filippo, Raúl A. Ortiz-Merino, Chiara Damiani, Gianni Frascotti, Danilo Porro, Kenneth H. Wolfe, Paola Branduardi, Dario Pescini

## Abstract

Genome-scale metabolic models are powerful tools to understand and engineer cellular systems facilitating their use as cell factories. This is especially true for microorganisms with known genome sequences from which nearly complete sets of enzymes and metabolic pathways are determined, or can be inferred. Yeasts are highly diverse eukaryotes whose metabolic traits have long been exploited in industry, and although many of their genome sequences are available, few genome-scale metabolic models have so far been produced. For the first time, we reconstructed the genome-scale metabolic model of the hybrid yeast Zygosaccharomyces parabailii, which is a member of the Z. bailii sensu lato clade notorious for stress-tolerance and therefore relevant to industry. The model comprises 3096 reactions, 2091 metabolites, and 2413 genes. Our own laboratory data were then used to establish a biomass synthesis reaction, and constrain the extracellular environment. Through constraint-based modeling, our model reproduces the co-consumption and catabolism of acetate and glucose posing it as a promising platform for understanding and exploiting the metabolic potential of Z. parabailii.

## 1 INTRODUCTION

Current genome sequencing technologies allow a fast and cheap overview into the genetic composition of virtually any organism, but connecting such genotypes to observed phenotypes remains a challenge. The reconstruction of genome-scale metabolic networks provide structured frameworks to represent the biochemical transformations within a target organism as complex genotype-phenotype relationships. Afterwards, different modeling approaches can be used to simulate, understand, predict and eventually control the behavior of such genome-scale metabolic networks. Flux Balance Analysis (FBA) is a widely used constraint-based modeling approach that relies on linear programming and the optimization of a given objective function (e.g., maximization of growth) for the determination of the metabolic model flux distribution [1, 2]. This approach is based on the assumption that organisms operate under a series of constraints limiting their possible functions, and leading to the definition of allowable cell phenotypes from defined metabolic networks [3].

In the last two decades, genome-wide reconstructions of metabolism have been produced for a plethora of model organisms, spanning from bacteria to higher eukaryotes [4]. The original versions of these models typically undergo incremental improvements. In particular, 10 different versions of the *Saccharomyces cerevisiae* metabolic network have been produced to date, by implementing cellular compartments, curated reactions, standard nomenclature, and even transcriptional regulation, as reviewed in [5]. However, less extensive efforts have been dedicated to other so-called non-conventional or non-*Saccharomyces* yeast species, despite their relevance for biotechnological applications, as well as for basic and biomedical research. The few non-conventional yeast species for which genome-scale metabolic reconstructions have been made to date are: *Pichia pastoris* [6, 7, 8, 9], commonly used as a host for recombinant protein production, *Scheffersomyces stipitis* [10, 11, 12], relevant for biomass bioconversion, the fission yeast *Schizosaccharomyces pombe*, especially relevant for cell cycle control studies [13], the opportunistic pathogens *Candida glabrata* and *C. tropicalis* [14, 15], used for drug target prediction, and lipid production, *Yarrowia lipolytica* [16, 17, 18, 19], relevant for its ability to produce and accumulate lipids potentially useful for biofuel production, and *Kluyveromyces lactis* [20, 21], remarkable for its characteristic lactose metabolism and its use for heterologous protein production. These models have however all been reconstructed starting form a reference *S. cerevisiae* model. Nevertheless, it is important to mention that yeast biodiversity is huge, and it goes beyond the models reconstructed up to now. In this regard, species in the *Z. bailii sensu lato* clade are known for their exceptional tolerance to different types of stress. In the case of *Z. bailii*, high tolerance to low pH and weak acids make these yeasts notorious as frequent agents of food spoilage [22, 23]. However, the same tolerance traits can also be seen as beneficial for the microbial-based production of otherwise aggressive compounds such as lactic acid [24].

Here, we present the first genome-scale model reconstruction and analysis of *Zygosaccharomyces parabailii*, which is one of the 3 species described in the *Z. bailii sensu lato* clade and one of the 12 formally described species in the *Zygosaccharomyces* genus [25, 26].

This work is based on our recent genome assembly and annotation, which we have improved by producing functional annotations and used to obtain transcriptional profiles upon lactic acid stress [27]. Our model does not necessarily reflects the hybrid nature of *Z. parabailii* ATCC60483, but it can be pivotal for making progress into the simulation and understanding of the high stress tolerance and metabolism of this yeast species, unveiling how to better exploit its nature for industrial purposes.

## 2 MATERIALS AND METHODS

### 2.1 Genome-scale metabolic model reconstruction

The reconstruction procedure consists of seven major steps:

#### 1. Annotate the genome of the target organism and highlight its metabolic functions

The *Z. parabailii* genes were automatically obtained with an improved version of the Yeast Genome Annotation Pipeline based on homology and synteny information from 20 different yeast species from the Saccharomycetaceae family [28]. These gene predictions were curated using Illumina RNAseq data using the Trinity / Pasa pipeline followed by manual inspection [29]. Afterwards, Enzyme Commission (EC) numbers were obtained using Blast2GO as previously reported [27, 30]. These EC numbers were inferred based on sequence homology keeping track of the organisms from which this information is taken.

#### 2. Populate a first draft model by connecting genes associated with an EC number into corresponding metabolic reactions

This was done with the Python package Bioservices [31], which was used to connect to the KEGG database [32, 33] to extract all the enzymatic reactions and related functional hierarchy information for every *Z. parabailii* gene with an assigned EC number. The KEGG functional hierarchy was used here to filter out all the EC numbers from enzymes whose function is not related to metabolism.

#### 3. Refine the draft model using information from phylogenetically close organisms for which genome-scale metabolic models are available

When possible, we inferred reaction compartmentalization, which is not represented in the KEGG database, and stoichiometry from the iOD907 model for *K. lactis* and the Yeast7 model for *S. cerevisiae*, taken here as references [20, 34]. This process is guided by the EC-homology information stored from step 1. Moreover, reactions from the reference models whose Gene-Protein-Reaction (GPR) rule is fulfilled are also included in the model draft. The GPR rules exploit a boolean expression for indicating which genes are involved in a given reaction and how they are interconnected. The satisfaction criterion implies that if a rule has the form “gene A AND gene B”, both genes in the reference model must be homologs of the draft model genes. On the contrary, if the rule has the boolean operator OR, just one of the two genes is necessary to be homolog. All the reactions from the reference models that are not associated to any GPR rule, excluding transport and exchange reactions, are *a priori* included into the draft model.

Special considerations were taken for reactions in the draft model involving genes whose EC numbers where not inferred either from *S. cerevisiae* or *K. lactis*. In these cases, the stoichiometry of the biochemical reaction was taken from the KEGG database, whereas the localization was assigned to the cytosol compartment unless otherwise stated in the Uniprot database [35]. Manual curation was performed for reactions in the KEGG database that are catalogued as “general reaction” involving generic substrates and products. For example, the KEGG reaction with identifier R01532 corresponds to the equation *Nucleoside triphosphate + H*_2_*O <=> Nucleotide + Diphosphate*. This reaction is labelled as general because its main substrates and products, namely Nucleoside triphosphate and Nucleotide, are generic compound that can correspond to ATP, CTP, UTP, GTP or TTP in case of Nucleoside triphosphate, and to AMP, CMP, UMP, GMP or UMP in case of Nucleotide. In these cases, the KEGG database suggests a group of specific reactions that are all included into the network under construction.

#### 4. Add transport reactions for metabolites present in multiple compartments

To our knowledge, transport reactions have not been fully characterized in *Z. parabailii*. Therefore, a reversible and unbounded transport reaction (i.e. lower and upper bound are set equal to −10^6^ and 10^6^ mmol gDW^−1^ h^−1^) is always added between the cytosol and any other compartment for every metabolite. For example, if a metabolite *m* is located in the cytosol (*m*_cyt_), in the mitochondrion (*m*_mit_) and in the peroxisome (*m*_per_), two transport reactions are added to the model, namely *m*_cyt_ ↔ *m*_mit_ and *m*_cyt_ ↔ *m*_per_. If a metabolite *m* is located in multiple compartments, such as, the mitochondrion (*m*_mit_) and the peroxisome (*m*_per_), but neither is in the cytosol compartment, the metabolite *m*_cyt_ is first added to the model and then two transport reactions are included, namely *m*_cyt_ ↔ *m*_mit_ and *m*_cyt_ ↔ *m*_per_.

#### 5. Integrate experimental chemostat cultivation and medium composition data

This step is optional as it depends on the availability of chemostat data and information about the medium composition. The data from chemostat cultivation is integrated into the model by adding an exchange reaction, which is a transport reaction between the intracellular and the extracellular environment for every metabolite whose production or consumption rate has been experimentally determined. Lower and upper bounds of these reactions are then constrained according to the measured values. Before integrating available experimental concentration values as exchange reaction constraints, these values were transformed into corresponding consumption or production rates *q_m_* using the as following equation:

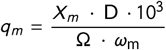

where *X_m_* is the concentration of the metabolite *m* measured in g l^-1^, D is the dilution rate measured in h^-1^, Ω is the dry cell weight measured in g l^-1^, and *ω*_m_ is the molecular weight of the metabolite *m* measured in g mol^-1^. A factor of 10^3^ was used to convert mol gDW^−1^ h^−1^ into mmol gDW^−1^ h^−1^, which is the standard measure unit of the FBA. The model can be further constrained by defining the set of exchange reactions from the extracellular environment in order to replicate the experimental growth medium by adapting its metabolite composition to the draft model. The lower bound of each exchange reaction 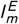 is set as follows:

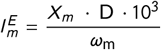

where *X_m_* is the concentration of the metabolite *m* within the experimental medium measured in g l^-1^, D is the dilution rate measured in h^-1^, and *ω*_m_ is the molecular weight of the metabolite *m* measured in g mol^-1^. As above, a factor 10^3^ is used to pass from mol gDW^−1^ h^−1^ to mmol gDW^−1^ h^−1^.

#### 6. Gap filling

Gaps are missing reactions leading to the formation of metabolites involved in just one reaction within the model as substrate or product, or in more than one reaction but only as substrate or product. These metabolites are generally called dead-end metabolites. The gap filling process consists of three steps:

- Dead-end metabolites are identified and catalogued as substrate or product, and their compartment localization is stored.
- The Bioservices package is used to automatically find in the KEGG database the reactions where each dead-end metabolite is involved
- Reactions involving dead-end metabolites as substrates are intersected with those involving dead-end metabolites as product to determine common identifiers. In case of a positive match, both metabolites are required to be localized into the same compartment while checking that the reaction was not already included into the model.

#### 7. Include exchange reactions to achieve mass-balance in the system

This last step involves a new search for dead-end metabolites for which an entry or exit irreversible exchange reaction is added to the draft model. If the unique reaction where a dead-end metabolite is involved is reversible, we added a reversible exchange reaction regardless of the role of the dead-end metabolite a substrate or product. The same criterion is followed for dead-end metabolites involved in more than one reaction: if all the involved reactions are irreversible, an exchange reaction is added to the model in the direction of uptake for substrate and secretion for product. When at least one of the involved reactions is reversible, an unbounded exchange reaction is included into the model.

### 2.2 Definition of the biomass synthesis reaction

The biomass synthesis reaction takes into account all the experimentally identified components where each element *m* is associated to a stoichiometric coefficient *s_m_* and computed as follows:

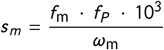

where *f*_m_ represents the fraction in weight of the monomer *m* into the macromolecule *P*; *f*_P_ represents the fraction in weight of the macromolecule *P* into the biomass, and *ω*_m_ is the molecular weight of the monomer *m*.

Glycerol, trehalose, chitin and 1,3-beta-D-glucan, take part in biomass formation according to the growth reaction composition of the *S. cerevisiae* and *K. lactis* genome-scale models. However, as information about weight percentages was only available for glycerol, we added to the model a mock reaction where carbohydrates contribute to the formation of a generic compound called “carbohydrates” that we included into the biomass reaction associated to a stoichiometric coefficient calculated similarly as above:

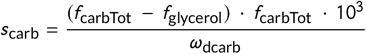

where *f*_carbTot_ is the fraction in weight of all the carbohydrates into biomass, *f*_glycerol_ is the fraction in weight of glycerol into carbohydrates, whereas *ω*_carb_ is the molecular weight of the fictitious “carbohydrates” species.

Since we had no information about single deoxynucleotide and nucleotide composition but we had their total percentages in weight, we added to the model two mock reactions for the formation of total DNA and RNA. In these mock reactions, the sum of the corresponding deoxynucleotides and nucleotides contribute to the formation of two generic compounds that we called “DNA” and “RNA”. These were included into the biomass reaction, and associated to a stoichiometric coefficient as follows:

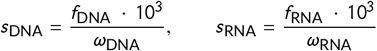

where *f*_DNA/RNA_ represents the fraction in weight of total DNA or RNA into biomass, and *ω*_DNA/RNA_ is the molecular weight of the fictitious DNA or RNA compounds.

As the total ATP cost necessary for the cell growth has not been experimentally determined, it was set to the value used in the biomass synthesis reaction of the *S. cerevisiae* and *K. lactis* genome-scale models.

### 2.3 Computation of the biomass yield

The biomass yield *Y* corresponds to the amount of biomass produced per grams of consumed carbon source, giving information about substrate conversion efficiency. This parameter is calculated as ratio of model simulations deriving flux of the biomass synthesis reaction over that of the glucose exchange reaction that is necessary to convert from mmol gDW^−1^ h^−1^ into grams. To sum up:

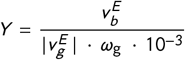

where 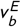 is the flux value of the biomass synthesis reaction, 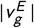 corresponds to the glucose exchange reaction flux value which is taken as absolute value to remove the negative sign due to exchange reactions convention, and *ω*_g_ is the glucose molecular weight.

### 2.4 Classic and parsimonious Flux Balance Analysis

The Flux Balance Analysis (FBA) is a constraint-based approach which exploits the linear programming to identify the optimal flux distribution that maximizes or minimizes a specific metabolic objective [1]. FBA relies on a steady state assumption, according to which time variation of each of the internal metabolites concentration is equal to zero. This means that *dX_i_* /*d t* = 0, where *X_i_* is the concentration of the metabolite *i*. The flux distribution represents the output of this analysis and shows the rate at which each reaction of the model occurs at steady state.

The application of constraints on the system under evaluation is necessary to reduce the set of candidate flux distributions, defining in this way an allowable solution space where any flux distribution may be equally acquired by the model. The maximization or minimization of a specific objective function *Z* allows to get a further narrowed feasible solutions space, and to identify a single optimal flux distribution.

The set of metabolic reactions included in the model is mathematically represented as a stoichiometric matrix *S* of size *M * R*, where *M* is the number of metabolites and *R* is the number of reactions included in the model. Given the stoichiometric matrix *S*, where each element *s_i_*_,*j*_ represents the stoichiometric coefficient of each metabolite within each reaction, the formulation of a FBA problem may be formulated in this way:

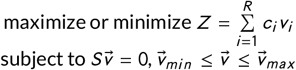

where *c_i_* indicates the objective coefficient value for the reaction *i*; *v_i_* represents the flux value of the reaction *i*; the two vectors 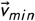 and 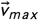 represent the lower and upper bounds vectors, expressing for each flux *v_i_* of the vector 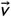, the range within it can vary. An interval between negative and positive values means that flux through the corresponding reaction can flow in the backward or in the forward direction.

The parsimonious FBA (pFBA) [36] is a variant of the classic FBA approach that applies the same principles, but it set as objective function the minimization of the sum of all fluxes. Therefore, pFBA assumes that there is a selection for the fastest growing strains that minimize the total necessary resources to implement the optimal solution.

Although the FBA only returns a single flux distribution, the constraints imposed on the system under investigation do not always allow to obtain a unique solution, but may confine the solution space to a feasible set of alternative optimal flux distributions in which the same optimal flux value of the objective function may be reached through different but equally possible ways. In this context, the Flux Variability Analysis (FVA) [37] returns the range of flux variability of each reaction, i.e. the allowable minimum and the maximum fluxes by each model reaction.

All the FBA and FVA simulations has been performed by using the COBRApy Toolbox functions [38] and the Gurobi solver.

### 2.5 Reaction deletion analysis

The gene deletion analysis is based on the knowledge for all reactions of the model under investigation of the corresponding gene-protein-reaction (GPR) rule, and implies that the deletion of each gene included in the model is cyclically simulated. This means that flux through all the reactions in which this gene is involved, according to the corresponding GPR rule, is blocked. The reaction deletion analysis is conceptually similar to the gene deletion analysis. It is performed by cyclically setting to zero both lower and upper bounds of each reaction in the model, preventing flux passing through it, without addressing the corresponding GPR rule. A FBA running is then performed to analyze the effect of each deletion on the objective function flux value.

### 2.6 Particle Swarm Optimization

The Particle Swarm Optimization (PSO) is a population-based stochastic algorithm, originating from studies of birds flocking, which aims at finding the set of parameters that maximize a specific objective [39]. The key concept behind the PSO is that an initial population of several candidate solutions, called particles, are randomly located inside a constrained search space by cooperating one with each other for finding the global best solution. Each candidate solution is then evaluated by the PSO algorithm by considering how much the particle minimizes its distance from the objective function, determining in this way the fitness value of that solution. Each particle, which is characterized by both a position in the search space and a velocity, keeps track of its personal best position that it has achieved so far in the search space. At each iteration of the algorithm, the velocity of the particle updates its current position, and the fitness is then revaluated. The individual best positions of each particle and the global best position achieved among all particles in the swarm are updated by comparing the current fitness values with the previous ones, and replacing them if better fitnesses are obtained. The PSO algorithm continues until stopping condition is reached. The final solution corresponds to the best fitness value achieved among all particles in the swarm, referred to as global best fitness, and the particle that achieved this fitness, referred to as global best candidate solution.

We implemented the PSO algorithm by using Python as wrapper. We carried out this analysis by setting a swarm size of 32 particles, 2000 iterations, and a range between 10^−6^ and 1 for the coefficients to estimate. We computed the fitness function by using the least square method to calculate the minimal distance between the computational and the experimental biomass yield values.

### 2.7 Experimental methods

#### Glucose-limited aerobic chemostat cultivations

Chemostat cultivations were performed in Biostat-B fermenters (B-Braun). A defined medium with vitamins and trace metals was used [40]. The glucose concentration in the reservoir medium was about 20 gl^−1^. A constant working volume of 1300 ml was maintained via an effluent line coupled to a peristaltic pump. A dissolved oxygen concentration above 50% of saturation was maintained by an air flow of 1.3 l^−1^min^−1^ (1 v/v/m) and a stirrer speed of 1000 rpm. The temperature was maintained at 30°C and the culture pH at 5.0 by automatic addition of 2 M KOH. The dilution rates were set at 0.1 h^−1^ (fully respiratory metabolism) or at 0.3 h^−1^ (respiro-fermentative metabolism). Cultures were assumed to be in steady state when at least seven volume changes had passed since the last change in growth conditions and the culture did not exhibit metabolic oscillations. The dry cell weight (Ω) was determined by filtering 10 ml of culture broth through pre-dried 0.22 *μ*m membranes filters. The filters were washed with demineralized water and dried to constant weight in a microwave oven [41]. For every steady state, during chemostat cultivation, the percentages of O_2_ and CO_2_, present in the off-gas were measured with a gas analyzer (Bioindustrie Mantovane, with Peltier system for O_2_ reading and infrared for CO_2_).

#### Analysis of extracellular metabolites

2 ml of culture broths were centrifuged for 2 minutes at maximum speed in microcentrifuge. Supernatants were stored at −20°C until analysis. At the time of the analysis, the samples were diluted with H2O milli-Q according to the need and loaded onto a HPLC apparatus (Jasco), equipped with UV (210 nm) and RI (refractive index) detectors. A BioRad Aminex HPX-87H column was used for metabolites separation. The column was maintained at 35°C, the flow at 0.5 mlmin^−1^ and the mobile phase was H_2_SO_4_ 0.005 N.

#### Lyophilization of samples for macromolecular biomass analysis

When the chemostat cultures reached the steady state of growth, an aliquot of culture broth of about 100 ml (a variable amount, needed to recover about 1 g of biomass) was collected by centrifugation at 5000 g for 7 minutes at 4°C; the supernatants were discarded and the pellets were resuspended in about 45 ml of 20 mM Tris-HCl (pH 7.6) each. Aliquots of 2 ml of sample were transferred to microfuge tubes and, after a second spin cycle of 2 minutes at maximum speed in microcentrifuge, the pellets were quickly frozen by immersion in a equilibrated bath of 50% ethanol and dry ice. The frozen samples were temporarily stored at −80°C and then loaded on the centrifugal evaporator Eppendorf Concentrator 5301 for about 1 h and 30 min. The lyophilized samples where stored at −20°C.

#### Biomass elemental and macromolecular analyses

The analyses were carried out using the lyophilized samples obtained from the chemostat cultures at dilution rates of 0.1 h^−1^ and 0.3 h^−1^. The biomass elemental analysis was performed in outsourcing by REDOX S.r.l. (Monza, MB, Italy).

For total protein determination, lyophilized biomasses were lysed before proceeding with the determination of the total proteins. The lysis efficiency has been verified by microscopic observation and by the Micro BCA (Thermo) colorimetric assay. Cells were collected as previously described and resuspended in lysis buffer (Tris-HCL 25 mM, pH 7.5 EDTA 25 mM) containing protease inhibitors (PMSF 0.5 mM and complete protease inhibitors, Roche). The samples were resuspended in 1 ml of lysis buffer and microfuge tubes were prepared, containing aliquots of 600 *μ*l of sample and 600 *μ*l of glass beads were distributed in microfuge tubes placed in a pre-chilled rack, loaded onto the TissueLyser II apparatus (Qiagen) and subjected to three stirring cycles at maximum power (30 Hz), lasting 5 minutes each. For the quantification of total proteins the Micro BCA assay (Thermo) was used. The lysate samples were diluted 1:1000 with H_2_O milli-Q and a 1 ml aliquots were added to 1 ml of reagent (prepared by mixing solutions A, B and C provided in a ratio of 25:24:1). The samples and the BSA standards, for the calibration curve set up, were incubated at 60°C for 1 h. After cooling, samples were transferred into cuvettes for spectrophotometric determination at 562 nm. The determination of the amino acid and lipid composition were performed in outsourcing by the laboratories of Analysis and Peptide Synthesis of the Centres Cientifics I Tecnològics of Barcelona (Spain)and by the Research Center on Metabolism (CEREMET) of the University of Barcelona (Spain), respectively. For total carbohydrates determination, the lyophilized pellets were resuspended in H_2_O milli-Q and diluted appropriately to obtain 1 ml aliquots containing about 0.1 mg of cells. Each sample was placed in a 15 ml tube with 5 ml of phenol 5% and 1 ml of H2SO4 96% and incubated at 90°C for 10 min. Solutions at increasing concentrations of glucose were similarly treated in order to then generate a calibration curve and quantify the carbohydrates present in the samples. Quantification was performed by spectrophotometric analysis at 488 nm. For the determination of glycogen content, samples were resuspended in 10 ml of 0.6 MHCl incubated 1 h at 100°C, then brought to room temperature, filtered with 0.22 *μ*m filters and subjected to an enzymatic assay (D-glucose–Hk enzymatic assays, Megazyme International Ireland). This assay measures the glucose released from the lysis of glycogen. For the total RNA assay the orcinol method was used [42]. The lyophilized pellets were resuspended in 3 ml of cold 5% trichloroacetic acid (TCA) and then centrifuged at 3500 rpm at 4 °C for 7 min, washed twice in the same solution. The residual pellets were left at −20°C for about 12 hours, then slowly thawed in melting ice and finally resuspended in 3 ml of perchloric acid (PCA) 0.3 M and placed at 90°C for 30 minutes. After cooling, the samples were centrifuged at 3500 rpm for 7 min and the supernatants were collected for RNA analysis. The reagent for the orcinol assay was assembled by dissolving 1 g of crystalline orcinol monohydrate and 0.5 g of FeCl hexahydrate in 100 ml of 37% HCl. The reaction mix was assembled in 15 ml glass tubes containing an aliquot of the sample (from 50 to 200 *μ*l), adjusted to 1.5 ml with H_2_O milli-Q and 1.5 ml of orcinol reagent. The tubes were gently stirred and placed at 90°C for 20 min, covered by glass beads to limit the evaporation. After cooling the samples, absorbance at 660 nm was measured at the spectrophotometer. The calibration curve was prepared with known concentrations of standard *S. cerevisiae* RNA. The total DNA was assayed using the Qubit apparatus (Invitrogen) and the kit associated with it HS-DNA. The lyophilized pellets were resuspended in 1 ml of lysis buffer, lysed by TissueLyser II (Qiagen), with the same methods described above and then diluted 1:1000 and 1:100 in the TNE buffer. The assay was carried out according to the supplier’s protocol.

## 3 RESULTS

### 3.1 Reconstruction of *Z. parabailii* metabolic model

We started the reconstruction process from the high-quality whole-genome sequence and annotation of *Z. parabailii* ATCC60483 [27]. We used the information in the KEGG database to filter out non-metabolic genes, and to extract the enzymatic reactions associated to EC numbers identified by our previous functional annotation [32, 29]. In this way, we selected 807 metabolic genes that were used to build the first draft of the *Z. parabailii* genome-scale matabolic network. We then obtained the stoichiometry and localization for the *Z. parabailii* reactions from the *S. cerevisiae* Yeast7 and the *K. lactis* iOD907 models when possible [20, 34]. Otherwise, the stoichiometry was derived from the KEGG database and the compartmentalization information was obtained from Uniprot or manually curated. Transport and exchange reactions were added to the draft metabolic network assuming a “matriosca”-like structure for the callular compartments. In this way, the extracellular environment represents the most external level containing the cytosol, which in turn includes all the other considered compartments of the model, which are mitochondrion, endoplasmic reticulum, lipid particle, cell envelope, nucleus, peroxisome, Golgi apparatus and vacuole. If a metabolite is available in multiple compartments, we added an exchange reaction only for the metabolite in the outermost compartment. Since no data are available for constraining these reactions, when the exchange reactions imply the consumption of a given metabolite from the extracellular environment, we set the lower bound to 5*10^−5^ mmol gDW^−1^ h^−1^ according to the coefficient inferred through the Particle Swarm Optimization (PSO) (see Section 2.6) to minimize the distance between the experimental and the computational biomass yield. The proposed reconstruction process is summarized in Figure 1.

**FIGURE 1.**
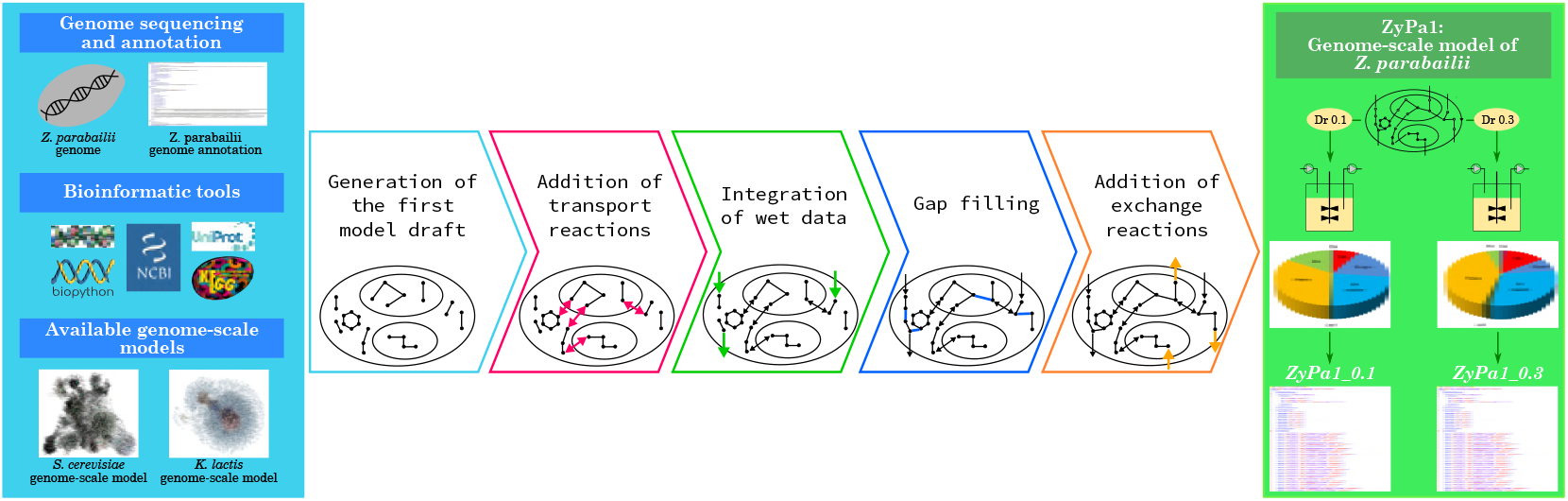
Graphical representation of the model reconstruction procedure.

Overall, the reconstruction process resulted in a model consisting of 3096 reactions (1743 of which are internal reactions, 619 are transport reactions and 734 are exchange reactions), 2091 metabolites and 2413 genes. Each reaction was associated to a GPR rule by joining the related genes with the OR boolean operator. Reactions and metabolites included in the model are localized over 10 different cellular compartments. The final version of the model, which we refer as ZyPa1, also includes the biomass synthesis reaction that we reconstructed according to the macromolecular biomass composition that we experimentally determined at the two dilution rates of 0.1 and 0.3 h^−1^. Because of the different stoichiometry in the two biomass synthesis reactions, we generated two versions of ZyPa1 model, one for each dilution rate, which we refer as ZyPa1_0.1 and ZyPa1_0.3, correspondingly.

We constrained the extracellular environment of the two ZyPa1 models according to experimental data. In particular, the exchange rates of glucose, oxygen, carbon dioxide, glycerol, acetic acid, acetoin and ethanol have been constrained based on the data obtained from chemostat cultivation at the two dilution rates of 0.1 and 0.3 h^−1^. They have been determined by scanning different dilution rates comprised in this interval. In Figure 2B, the experimentally determined metabolic profiles are given. A second set of constraints on the extracellular environment concerned the *in silico* medium composition in order to replicate the growth on the experimental medium, by adapting it to the model composition. Therefore, we integrated an exchange reaction for ammonium, biotin, (R)-pantothenate, nicotinate, myo-Inositol, thiamine, pyridoxal, 4-aminobenzoate and iron.

**FIGURE 2.**
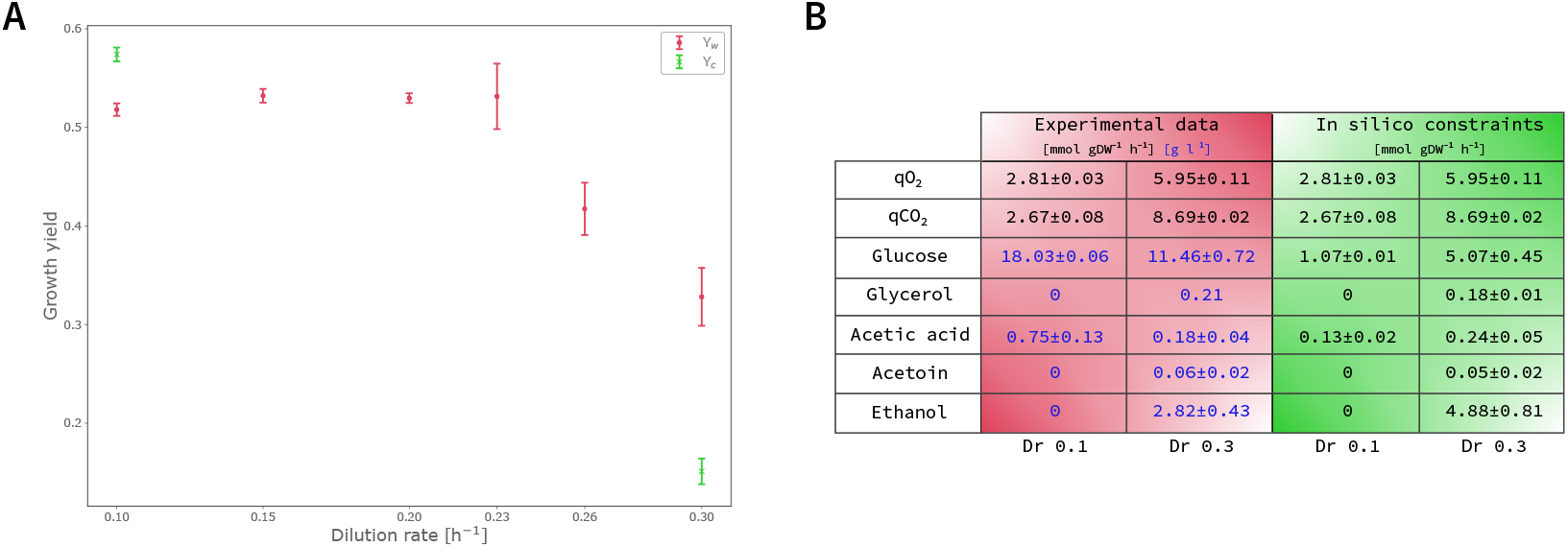
Comparison between experimental and computational data. (A) Computation of biomass yield at different dilution rates. Red circle points show the experimental biomass yield (*Y_w_*) computed at the six dilution rates of 0.10, 0.15, 0.20, 0.23, 0.26 and 0.30 h^−1^. Error bars for each point have been calculated according to experimental errors. Green x markers show the computational biomass yield (*Y_c_*) computed at the two dilution rates of 0.10 and 0.30 h^−1^. Error bars associated to each point is calculated according to glucose uptake flux variability. (B) Experimental data from chemostat cultivation at the two dilution rates of 0.10 and 0.30 h^−1^ are reported in the red part of the table. Different color are used to distinguish values that are measured in mmol gDW^−1^ h^−1^ (black) from values that are measured in g l^−1^ (blue). Computational constraints imposed on the two ZyPa1 models are reported in the green part of the table. All values are expressed as mmol gDW^−1^ h^−1^.

We checked for the presence of blocked reactions into the two ZyPa1 models in order to find all points in the network through which no flux pass, meaning that, following a FVA simulation, minimum and maximum flux of these reactions are equal to zero when no objective function is set. Currently, 75 and 72 reactions, *i.e.* about 2% of the total number of reactions, emerged as blocked respectively in the ZyPa1_0.1 and ZyPa1_0.3 models, with the only difference between them regarding the three exchange reactions of glycerol, acetoin and ethanol that we imposed equal to zero in ZyPa1_0.1 model according to the experimental data. The identified blocked reactions are caused by gaps of knowledge in the metabolic pathways of the target organism. However, since a genome-scale metabolic model, as such, integrates all the knowledge about metabolism of a given organism, these reactions are integral part of the model itself even if they are not, currently, able to carry flux.

The ZyPa1 model was deposited in BioModels [43] and assigned the identifier MODEL1807110001.

### 3.2 Comparison of model with other genome-scale models

We compared the ZyPa1 model with the genome-scale models used as reference during the reconstruction process, *i.e.* Yeast7 and iOD907, in order to highlight similarities and differences among the three networks. First, we calculated the size of the three networks in terms of involved reactions, metabolites and genes. As can be seen from Figure 3A, the number of reactions and metabolites that are included in the ZyPa1 model is closer to the corresponding value in *S. cerevisiae* than in *K. lactis*. However, it is worth to mention that the number of genes assigned to ZyPa1 network doubles that of Yeast7 and iOD907 models. This discrepancy is partially due to the interspecies hybrid nature of *Z. parabailii* as this organism has two copies of most genes, one deriving from each of the two independent parental genomes. In addition, some gene expansions occurred in the *Z. parabailii* genome. For example, in haploid *S. cerevisiae* cells, 3 genes encoding for the pyruvate decarboxylase activity, named *PDC1*, *PDC5* and *PDC6* have been annotated and described. Due to the hybrid nature of *Z. parabailii*, it was reasonable to expect to find 6 PDC genes, but the observed number of 12 suggests that in some cases expansion of genes number could justify the value we identified.

**FIGURE 3.**
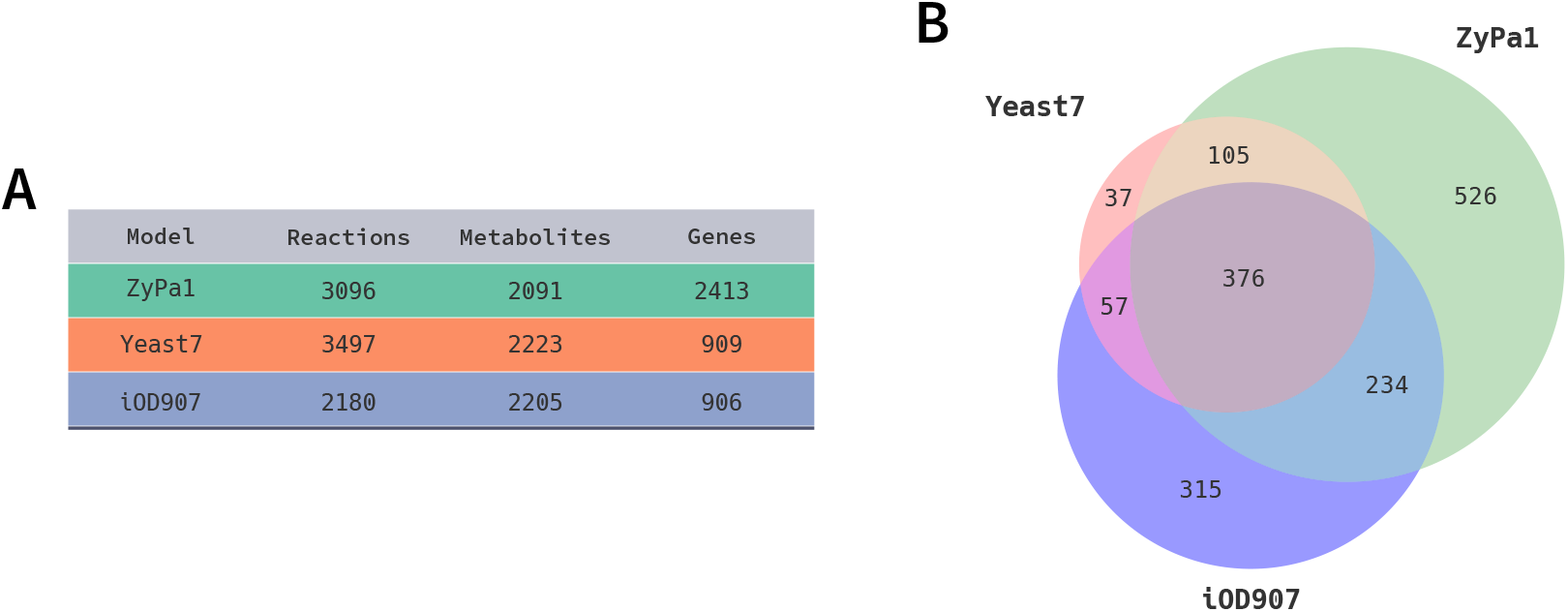
Comparison of ZyPa1 model with genome-scale metabolic models of *S. cerevisiae* and *K. lactis*. (A) Overview of the reactions, metabolites, and genes in the three models. (B) Venn diagram showing the overlapping reactions among the three models by using the related KEGG identifier. An unknown portion for each of the three network emerged for reactions having a missing reference to the KEGG database.

We extended the comparison among the three networks also at the level of the included reactions, and of pathways where they are involved. In Figure 3B, a Venn diagram shows the overlapping reactions among the three models by using, when available, the related KEGG identifier. An unknown portion for each of the three network emerged for reactions having a missing reference to the KEGG database. This portion is equal to 72% in Yeast7, 50% in iOD907 and 52% in our ZyPa1 model. By performing the comparison only on the known part of each network, we observed that ZyPa1 model does not rely as much on Yeast7, but rather on iOD907 model since the overlapping with *K. lactis* model is much higher compared to that with *S. cerevisiae* network.

For the set of reactions with a KEGG identifier, we collected all the metabolic pathways where they are involved. In Figure 4, the circle plot shows internally the hierarchy of all KEGG pathways related to metabolism. Each dead-end node corresponds to a specific pathway and the differently colored sections are needed for highlighting pools of pathways that are related one with each other. Around the outer circle, the relative frequency for each single pathway in the three genome-scale models over the total reactions number of the models themselves is shown. It is possible to observe variability among the three networks in terms of pathways distribution, and for some group of pathways the differences are more pronounced. This is the case of the group labeled as "LipidMetabolism". Focusing the attention on the comparison between *S. cerevisiae* and *Z. parabailii*, it is relevant to report that differences in phospholipids composition were proposed to be responsible of the different acetic acid membrane permeability in the two yeasts. More in detail, sphingolipids have been determined to be several times higher in *Z. bailii* than in *S. cerevisiae* and it was shown that upon acetic acid stress the membrane remodeling is more important in the non-*Saccharomyces* yeast and further exacerbates this difference [44]. By looking at the map, it does not seem to evidence a difference in the abundance of genes in the specific category of sphingolipids. Nevertheless, differences can be observed for other lipid categories: this can indicate a diverse ability of the two yeasts to remodel the lipid composition. Indeed, in a different study [45] it was demonstrated that lactic acid stress did not evoke important variations of the sphingolipid content of *Z. parabailii* cells, but a significant decrease in lipid acyl chain length occurring in the stationary phase of growth. This modification could contribute to lower membrane fluidity of organic acids in their undissociated form.

**FIGURE 4.**
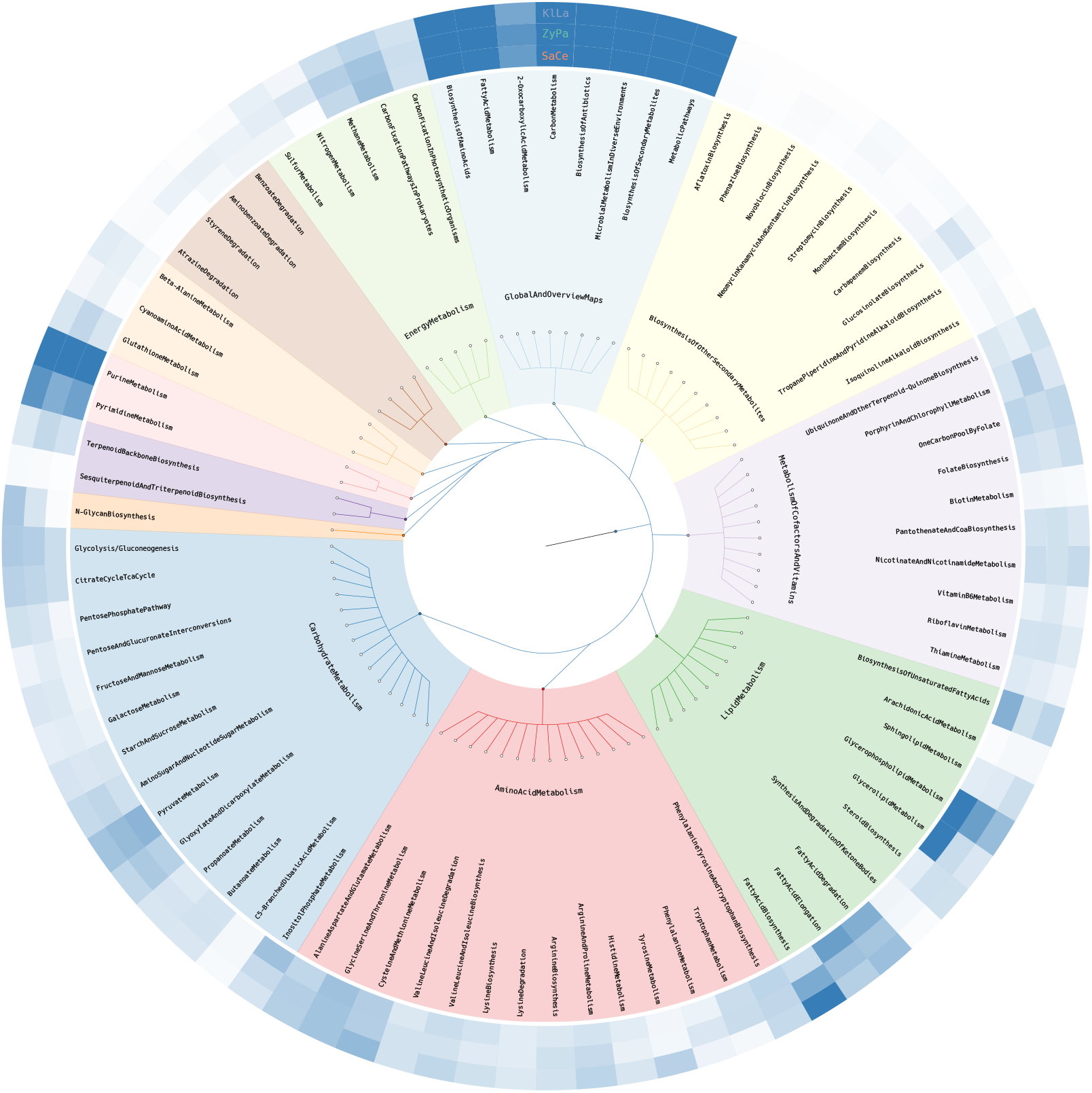
Metabolic pathways distribution among ZyPa1, Yeast7 and iOD907 genome-scale models. The circle plot shows the hierarchy of all KEGG metabolism related pathways included in the three models. Differently colored sections indicates pathways that are related one with each other. Each dead-end node corresponds to a specific pathway and the three outer concentric circles, starting from the innermost level, show the corresponding relative frequency in the genome-scale models of *S. cerevisiae* (SaCe), *Z. parabailii* (ZyPa) and *K. lactis* (KlLa).

### 3.3 Phenotype prediction

We assessed the in silico metabolic capabilities of the two ZyPa1_0.1 and ZyPa1_0.3 models by carrying out FBA simulations, and setting biomass synthesis reaction as the objective function to be maximized. The main readout resulting from the FBA is the calculation of the biomass yield that we obtained as ratio between the biomass synthesis flux value and the glucose exchange reaction flux value converted to grams.

From the FBA simulations, we computed the yield value for both ZyPa1 models and we compared them with the corresponding experimental values. As shown in Figure 2A, respectively at the dilution rate of 0.1 and 0.3 h^−1^, a biomass yield of 0.57 compared to the corresponding experimental value of 0.52, and a biomass yield of 0.15 compared to the corresponding experimental value of 0.33 emerged revealing good agreement of the simulations with experimental data. The computational biomass yield was calculated by taking into account the glucose uptake variability resulted from the FVA simulations indicated by the error bars in the plot.

### 3.4 Reaction deletion analysis

We performed a reaction deletion analysis of both ZyPa1 models to investigate which deletions have an effect on the biomass synthesis reaction flux value. It is worth noticing the impact caused by the in silico deletion of the R01290 reaction, which is catalyzed by the L-serine hydro-lyase enzyme and corresponds to the addition of L-Homocysteine to L-Serine, with the release of L-Cystathionine and one water molecule: L-Serine + L-Homocysteine <=> L-Cystanthionine + H_2_O. The simulation in both ZyPa1 models of R01290-catalyzing enzyme deletion implied a different reduction of biomass synthesis flux, in particular, of 23% in ZyPa1_0.1 model and a 39% in ZyPa1_0.3 model. This reaction is directly connected to the synthesis and metabolism of L-Cysteine, L-Methionine, along with metabolism of L-Serine, as substrate of R01290 reaction.

The small discrepancy between the stoichiometric coefficients associated to L-Cysteine, L-Methionine and L-Serine into the biomass synthesis reaction of both ZyPa1 models does not justify the observed reduction of biomass production flux value resulted from the knock out of R01290 reaction. A possible explanation for the result of this perturbation can be related to the ability of *Z. parabailii* to produce various aliphatic and aromatic alcohols known as fusel alcohols [46]. During food fermentation in yeast, fusel alcohols are produced from amino acid catabolism by using the so called Ehrlich pathway. A first transamination reaction generates the *α*-keto acid that is then decarboxylated to the corresponding aldehyde by *α*-keto acid decarboxylases. Several pyruvate decarboxylases (PDC) enzymes, among which those that generally participate in alcoholic fermentation by converting pyruvate to acetaldehyde can account for the reaction. A higher copy number of PDC genes in *Z. bailii* than in *S. cerevisiae* genome together with a less efficient Crabtree effect suggests that PDC genes in *Z. bailii* may be beneficial for the conversion of amino acids and reducing sugars to aldehydes by the Ehrlich pathway rather than for favoring alcoholic fermentation in this organism. Aldehydes are subsequently converted to higher alcohols (the fusel alcohols) and acids by, respectively, alcohol dehydrogenases and aldehyde dehydrogenases. The fewer copy number of genes encoding alcohol dehydrogenases in *Z. bailii* compared to *S. cerevisiae* could be the reason why it produces more aldehydes and less alcohols compared with S. cerevisiae. This route may represents a rewiring, also partial, of metabolism when sugars source is depleted and amino acids can be then used as energy and carbon sources. Our finding that *in silico* deletion of R01290 reaction causes a higher reduction of biomass synthesis flux value at dilution rate 0.3 than 0.1 h^−1^ when the metabolism is respiro-fermentative is in line with the usage of Erlich pathway during fermentation, but it needs to be validated through appropriate *in vivo* experiments.

### 3.5 Ability of *Z. parabailii* to consume acetate

*Z. parabailii* is highly tolerant to organic acids. Moreover, its ability to consume organic acids also in the presence of glucose has been described [47]. This ability to consume acetate in the presence of glucose and oxygen is of particular interest as acetate is released from pretreated lignocellulosic biomass and acts as an inhibitory compound for most microbial cell factories but not for *Z. parabailii*.

We assessed the ability of *Z. parabailii* to simulate this behavior by identifying in the model all the reactions where acetate is involved as substrate or product. Then, we carried out a parsimonious FBA (pFBA) for analyzing if flux of these identified reactions is changed when acetate is consumed from the extracellular environment. In this simulation, we did not force an uptake of acetate in the models, but we just gave them the possibility to consume this carbon source from the external environment. This means that we fixed the upper bound of acetate exchange reaction to zero and we cyclically increased its lower bound from 0 to a value of −10 mmol gDW^−1^ h^−1^, where the negative value is a convention used for the exchange reactions indicating a consumption of a given metabolite from the extracellular compartment into the model. We choose to perform pFBA because in order to investigate the organic acid tolerance behaviour of *Z. parabailii*, this approach returns the flux distribution that minimize the total flux of sources for reaching a given objective. Moreover, the solution that of the classic FBA involved a null acetate uptake flux value that is however possible according to FVA output.

From this analysis, we firstly checked if an uptake of acetate takes place in *Z. parabailii* models, and we then verified if biomass synthesis or the acetate involving reactions change their flux value according to an uptake of this metabolite. We observed that the yield of both versions of the ZyPa1 model do not vary when this carbon source enters the cell. This result suggested two alternative hypotheses. The first one is that in both Zypa1 models acetate is not being consumed from the extracellular environment, leading to an unchanged flux distribution. According to the second hypothesis, acetate is entering the cell but without contributing to growth. By analyzing flux through acetate exchange reaction and the 21 model reactions where acetate is involved, we are more prone to consider the second hypothesis as true. Indeed, following the FVA, we observed that a consumption of acetate is possible in both models.

We chose to investigate the flux distribution where the acetate uptake flux was the highest value for each tested intake level between 0 and 10 mmol gDW^−1^ h^−1^. Given the highly tolerance to organic acids in the presence of glucose, the Figure 5 briefly shows the catabolism of these two metabolites. Once acetate is consumed, we observed that it contributes to both the cytosolic and mitochondrial Acetyl-CoA (AcCoA) pool, which is involved in multiple pathways, including the fatty acids biosynthesis, the metabolism of some amino acids and the Krebs cycle. At the experimental condition of pH5, acetic acid is about 50% in its indissociated form. Therefore, it enters the cell mainly by simple diffusion: the higher intracellular pH (pH_*i*_) determines its deprotonation, causing negative effects both for acidification and for the negative effects of the counteranion. The *in silico* evidence of the acetate consumption without a significant impact on the biomass yield suggested that the catabolism of this organic acid can contribute to its detoxification, possibly providing ATP for the energy-consuming proton extrusion and 2C units for the remodeling of membrane lipids (see above).

**FIGURE 5.**
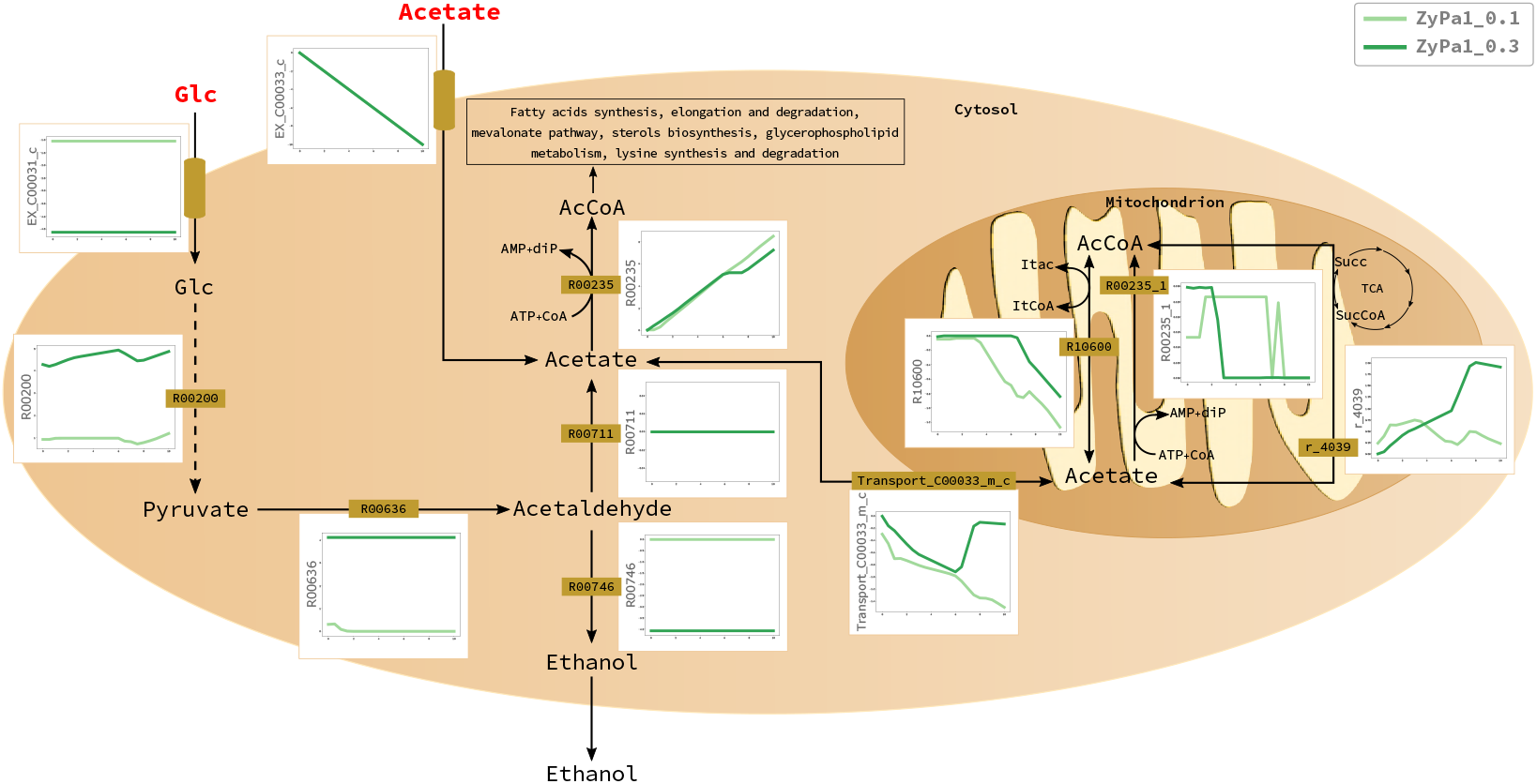
Effect of increasing acetate consumption rate from the extracellular environment between 0 and 10 mmol gDW^−1^ h^−1^ on ZyPa1 models flux distributions. Only the acetate involving reactions sensitive to acetate intake, and representative reactions from glucose catabolism are reported. In each plot, the x-axis shows the acetate uptake flux value, and y-axis shows the flux value for the reaction indicated along the axis. The light and dark green lines refer, respectively, to ZyPa1_0.1 and ZyPa1_0.3 models. Plots with a negative slope of the lines and, consequently, a negative flux value along the y-axis for the related reaction indicates that the reverse direction of the reaction is followed.

## DISCUSSION

In this work, for the first time we reconstructed the metabolic model of the stress tolerant hybrid yeast *Z. parabailii* ATCC60483 based on the recent genome assembly and annotation coupled with wet data obtained with chemostat cultivation at two different steady states, one fully respiratory and the second respire-fermentative. The two steady states were defined in terms of principal metabolites and macromolecular composition of the biomass.

This first version of the model proved to be able to describe the experimental biomass yield at a quite good level, and through the reaction deletion analysis revealed a different impact in terms of biomass synthesis flux reduction caused by the *in silico* deletion of the reaction catalyzed by the L-serine hydro-lyase enzyme. This reaction, which is connected to the metabolism of L-Cysteine, L-Methionine and L-Serine, may represent a metabolic rewiring when source of sugars is depleted and amino acids can be used as energy and carbon sources. It is worth noticing that when GPR rules associated to reactions in the model will be cured, it will be possible to run different simulations, among which the gene deletion analysis, which exploits the knowledge of genes involved in a specific reaction and their relationships. This will confer to the model the capability of prediction about knock out of specific genes in order to perform certain tasks.

This first version of ZyPa1 model also showed the ability to follow the acetate catabolism at different dilution rates. The *in silico* evidence for an acetate consumption without a significant impact on the biomass yield supports the hypothesis that the main contribution of this catabolism is in the detoxification of this organic acid as an inhibitory compound often released from pretreated lignocellulosic biomass.

It is worth to mention that the majority of the processes related to the production of bulk chemicals are run in batch or fed-batch and it is important also in this case to use wet data for asking the model for predictions. In this regard, the reconstructed genome-scale model can be used as a scaffold for implementing a reduced version of the network that can focus the attention on specific small-scale pathways. In this context, the investigation of the system dynamics can be performed by means of the mechanism-based modeling approach giving a detailed description of specific cellular processes with also the greatest predictive capability about the functioning of biological systems at the molecular level.

The differential analysis of gene abundance for diverse categories of functions can inspire novel experiments and help in the description of the peculiarity of this yeast.

## ACKNOWLEDGEMENTS

The institutional financial support to SYSBIO Center of Systems Biology - within the Italian Roadmap for ESFRI Research Infrastructures - is gratefully acknowledged. M.D.F. is supported by a SYSBIO fellowship. This work was also funded by the European Union FP7 Marie Curie Programme [YEASTCELL - 7PQ MARIE CURIE (12-4-2001100-40)] to PB and KW, supporting the PhD fellowship of ROM.

## AUTHOR CONTRIBUTIONS

MDF and DPE implemented the model reconstruction procedure of ZyPa1 model and performed all computational analyses. CD, PB and ROM assisted in data analysis. GF and PB generated all experimental data. PB and DPE supervised the project. MDF, DPE, PB, CD and ROM wrote the manuscript, which was reviewed by all authors.

